# A compact multifunctional model of the rabbit atrioventricular node with dual pathways

**DOI:** 10.1101/2023.01.30.526208

**Authors:** Maxim Ryzhii, Elena Ryzhii

## Abstract

The atrioventricular node (AVN) is considered a “black box”, and the functioning of its dual pathways remains controversial and not fully understood. In contrast to numerous clinical studies, there are only a few mathematical models of the node. In this paper, we present a compact, computationally lightweight multi-functional rabbit AVN model based on the Aliev-Panfilov two-variable cardiac cell model. The AVN model includes fast (FP) and slow (SP) pathways, primary pacemaking in the sinoatrial node, subsidiary pacemaking in the SP, and takes into account the asymmetry of coupling between model cells. At the same time, the model is accompanied by a visualization of electrical conduction in the AVN, revealing the interaction between SP and FP in the form of ladder diagrams. The AVN model demonstrates broad functionality, including normal sinus rhythm, AVN automaticity, filtering of high-rate atrial rhythms during atrial fibrillation and atrial flutter with Wenckebach periodicity, direction-dependent properties, and realistic anterograde and retrograde conduction curves in the control case and the cases of FP and SP ablation. To show the validity of the proposed model, we compare the simulation results with the available experimental data. Despite its simplicity, the proposed model can be used both as a stand-alone module and as a part of complex three-dimensional atrial or whole heart simulation systems and can help to understand some puzzling functions of AVN.

## 1 Introduction

The atrioventricular node (AVN) plays a key role in the cardiac electrical conduction system. It is located between the atria and ventricles, namely in the base of the right atrium (Fig 1A). The AVN is the only site responsible for transmitting impulses originating in the sinoatrial node (SN) to the ventricles of the heart, coordinating the relationship between contraction periods of the heart chambers, and introducing a variable delay, allowing effective pumping of the blood in a wide range of cardiac rhythms^1, 2^. Fig 1B demonstrates the detailed anatomical location of the rabbit AVN and surrounding tissues^3^.

**Figure 1.**
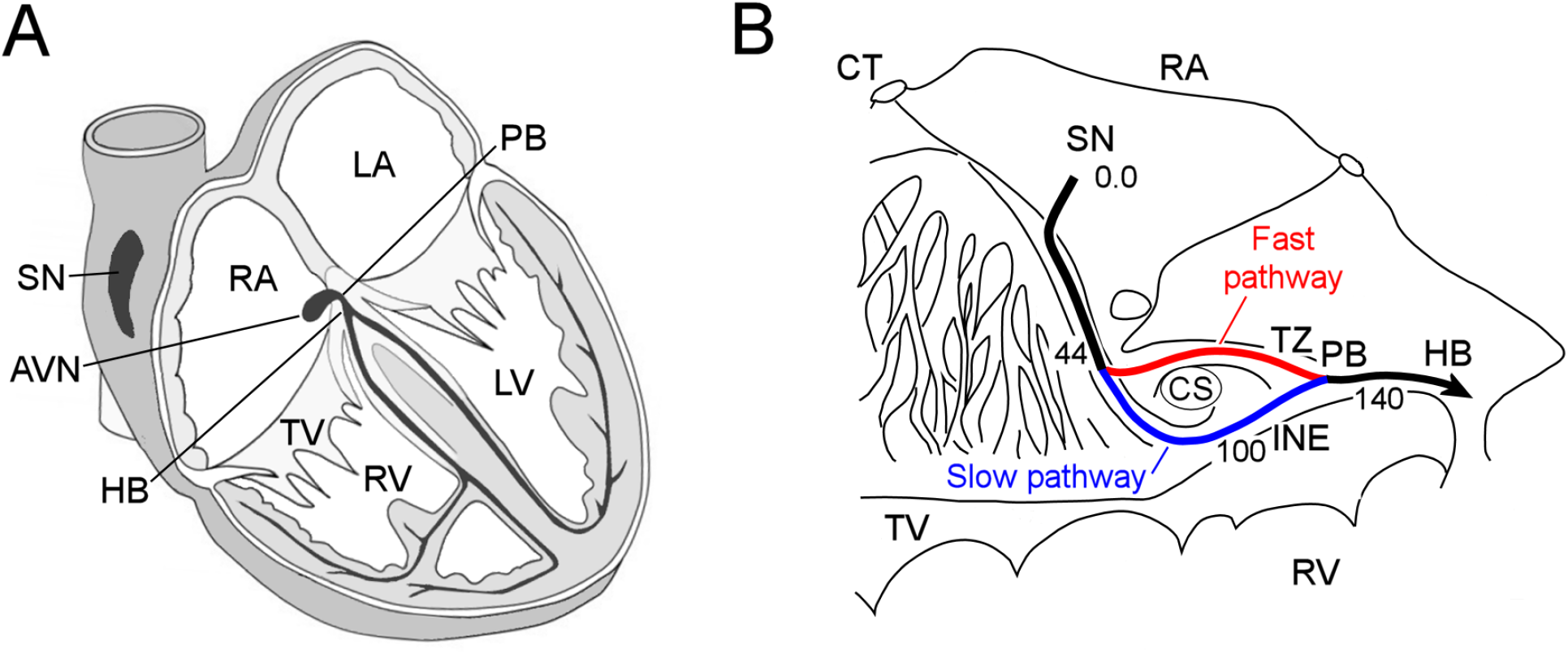
**(A)** Conduction system of the mammalian heart. SN - sinus node, AVN - atrioventricular node, RA and LA - right and left atria, PB - penetrating bundle, HB - His bundle, TV - tricuspid valve, RV and LV - right and left ventricles. **(B)**Anatomic drawing of the rabbit atrioventricular node and surrounding tissues with excitation wave latency (from^3^). CT - crista terminalis, CS - coronary sinus, TZ - transitional zone, and INE - inferior nodal extension. The numbers in milliseconds are given with respect to the sinus node pacemaker. The fast and slow pathways are indicated by red and blue lines, respectively.

The rabbit AVN contains two built-in functional pathways conducting impulses from the atria to His bundle (HB) - so-called fast pathway (FP) with a longer effective refractory period, and slow pathway (SP) with a shorter one (Fig 1B). The FP also has a smaller conduction latency than the SP^4–6^. The dual pathway AVN physiology provides protection of ventricles from atrial fibrillation (AFB) and supraventricular tachycardia by filtering excessive atrial electrical impulses^7^. On the other hand, the interplay of the FP and SP in the AVN can be a potential substrate for atrioventricular nodal re-entrant tachycardia (AVNRT)^6, 8^. The AVN also acts as a subsidiary pacemaker and provides atrial and ventricular pacing in the case of SN failure. According to experimental studies, in such a case, the AVN pacemaker activity in the rabbit heart originates in the SP between the coronary sinus and the tricuspid valve in the inferior nodal extension^9^.

Despite numerous research efforts, many aspects of the AVN electrophysiological behavior remain unclear and debated^1^. In addition to animal experiments and clinical studies on humans, mathematical modeling at different levels of detalization and complexity may offer insight into the AVN “black box”. Currently, very few AVN models or atrial models incorporating AVN with dual conduction pathway exist^10–15^. The rabbit cardiac biophysically-detailed multicellular model^10^ is complex and requires significant computational resources. The simulation of different AVN phenomena with this model requires reconfiguration, e.g., changing the number of model cells. Moreover, some reproducibility issues were reported recently^16^. The model computes normal action potential propagation from the SN through the atrium to AVN, AVNRT, the effect of the FP and SP ablation, *I_Ca,L_* block effect, the AVN filtering function during AFB, and AVN pacemaking.

There were also attempts to develop AVN models based on more computationally effective simplified nonlinear two-variable models. In the work^13^ the AVN model based on hundreds of Lienard-transformed modified Van der Pol^17^ and FitzHugh-Nagumo^18, 19^ equations, reproduces normal sinus rhythm and AVNRT. The three-dimensional anatomically-detailed model of the rabbit right atrium^14^ includes the AVN with dual pathways and utilizes the cellular automaton method and modified FitzHugh-Nagumo and Rogers^20^ two-variable models. The model computes normal sinus rhythm and nodal behavior during AFB. In work^21^ the human real-time hybrid automaton model was proposed, which can emulate sinus rhythm and the AVNRT. In the works^11, 12^ the authors used a simple, functional mathematical model of the rabbit AVN with the inclusion of dual pathway physiology. This model consists of exponential mathematical equations and is based on rabbit heart preparation data, and is able to predict AVN conduction time and the interaction between the FP and SP wavefronts during regular and irregular atrial rhythms. The functions of the models include simulation of nodal recovery, concealed conduction, Wenckebach phenomena, AFB and atrial flutter (AFL), and FP and SP ablations. The network model of human AVN^22^ consists of FP and SP interconnected at one end, which includes functions of the HB and Purkinje network. Both pathways are composed of interacting nodes, and the refractory periods and conduction delays of the latter are determined by the stimulation history of each node and described by exponential functions^23^. The model is able to represent RR interval series extracted from ECG data (both in the forms of histograms, Poincaré plots, and autocorrelation), and the latest version also incorporates autonomic tone^24^.

In our previous work^15^, we considered a simplified human AVN model with dual pathways, which consists of about 40 cells including excitable cells and pacemaking cells described by Aliev-Panfilov^25^ (APm) and modified Van der Pol models. The model reproduces normal sinus rhythm, AVN automaticity, reentry, and filtering function. However, we found that various variants of the modified Van der Pol equation are not suitable for the description of cells of subsidiary pacemakers.

In this work, we present a compact mathematical model of the rabbit AVN based on a single nonlinear two-variable APm for the description of both quiescent excitable and pacemaking cells, which allows representing various parts of cardiac conduction system with different tissue types in a uniform and convenient way. The model consists of only 33 cells and requires relatively low computational resources. Adjusting to available experimental data for the rabbit heart made it possible to demonstrate a wide range of behavioral phenomena of AVN. This includes proper bidirectional conduction in the FP and SP both in the normal case and after the FP and SP ablations reflected in the calculated recovery curves, AVNRT, the AVN automaticity, filtering function during AFB and AFL, and Wenckebach periodicity. Correct reproduction of retrograde conduction characteristics was achieved by introducing coupling asymmetry between model cells. In addition, we included visualization demonstrating the inner processes in the AVN “black box”, in particular, the laddergrams (Lewis ladder diagrams)^26^ with the exact timing of each model cell in both FP and SP pathways during the excitation wave propagation. Finally, the comparison with published experimental findings provides validation of the simulated results.

## Methods

### 1.1 Structure and Equations of the Model

A schematic view of the one-dimensional rabbit cardiac conduction system model incorporating AVN with dual pathways is shown in Fig 2. The model consists of a two-row matrix, where the upper row includes the SN and peripheral SN (PS1–PS3) cells, atrial muscle (AM1–AM3), fast pathway (FP1–FP8), penetrating bundle (PB), and HB (HB1-HB6) cells, while the lower row includes only ten slow pathway (SP1–SP10) cells and additional intermediate atrial muscle cell (AM*). The latter is added to provide a smooth transformation of short rectangular-shaped excitation impulses into typical atrial action potential (AP) shapes in the case of AFB (with random pacing) and AFL (regular pacing) simulations. The arrows *Antero* and *Retro* denote the points of application of external S1 and S2 stimuli for anterograde and retrograde propagation study with S1S2 stimulation protocol, and the arrow *AF* indicates entrance for “internal” stimuli in the cases of AFB and AFL simulations. The premature S1S2 stimulation^27^ refers to pacing that is performed at a constant basic cycle length (shorter than normal sinus rhythm) for 8–10 S1 impulses, followed by the introduction of an extra stimulus S2, and with stepwise reduction of the S1S2 interval until AVN refractoriness (total conduction block) is reached. Therefore, S1 and S2 mark the basic and test stimulus, respectively, and the same subscripts apply to atrial (A) and HB (H) stimulation or response.

**Figure 2.**
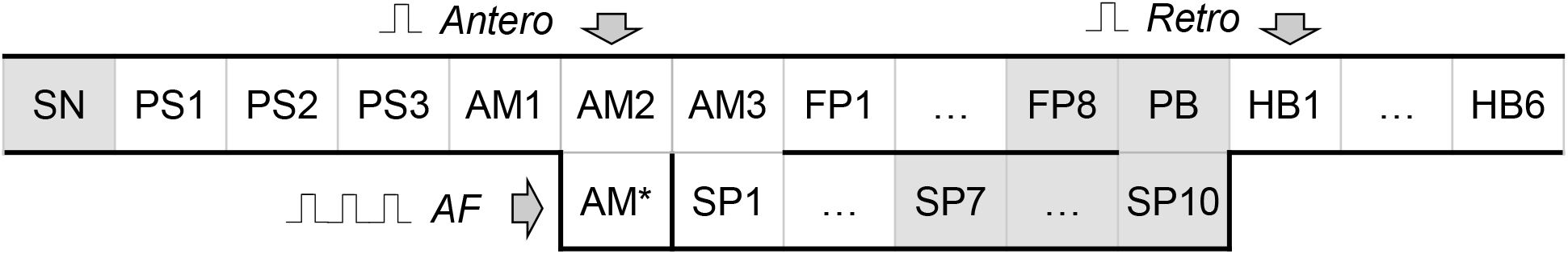
Schematic representation of the rabbit cardiac conduction model with dual pathway. SN - sinus node cell, PS1–PS3 - peripheral sinus node cells, AM1–AM3 - right atrial muscle cells, FP1–FP8 - fast pathway cells, SP1–SP10 - slow pathway cells, PB - penetrating bundle cell, HB1–HB6 - His bundle cells. Arrows *Antero* and *Retro* denote the points for anterograde and retrograde S1S2 stimuli application, arrow *AF* indicates the point of stimuli application for random pacing (atrial fibrillation) and regular pacing (atrial flutter), and AM* is an additional intermediate atrial muscle cell. The gray-shaded cells indicate primary and subsidiary pacemaker cells.

Each model cell represents a group of real cardiac cells and is described by two-variable APm^25^. The APm for monodomain description of the cardiac cell consists of the following set of reaction-diffusion type nonlinear ordinary differential equations:

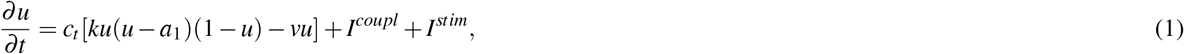

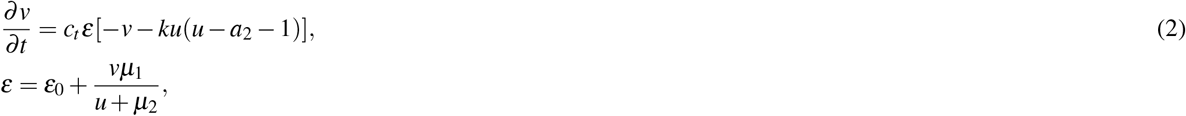

where 0 ≤ *u* ≤ 1 and *v* are the normalized transmembrane potential and slow recovery variables, respectively. Parameter *k* controls the magnitude of transmembrane current, and parameters *μ*_1_, *μ*_2_, *b*, and *a* are adjusted to reproduce characteristics of cardiac tissue, *ε* sets the time scale of the recovery process determining the restitution properties of the action potential, *c_t_* is the time scaling coefficient introducing physical time in seconds into the system. In the general case, *a*_1_ = *a*_2_ > 0. Since the APm was originally developed for the description of canine ventricular myocyte, the parameters were adjusted for the rabbit AVN in terms of action potential duration and refractoriness (see Supplementary Tables S1 and S2). *I^stim^* is the external stimulation current, *I^coupl^* = ∇ · (**D**∇*u*) is the coupling current, where ∇ is a spatial gradient operator defined within the model tissue geometry, and **D** is a tensor of diffusion coefficients characterizing electrotonic interactions between neighboring cells via gap junctional coupling conductance. For the considered one-dimensional system, the coupling term in the discrete form reads as follows

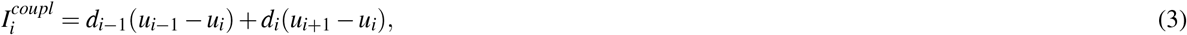

where *i* is the model cell index, and *d_i_*, is the diffusion coefficient in s^−1^ assigned for each pair of cells normalized on dimensionless distance. As will be discussed below, to enhance the direction-dependent conduction properties of the AVN model, we implemented a coupling asymmetry with the modified coupling term:

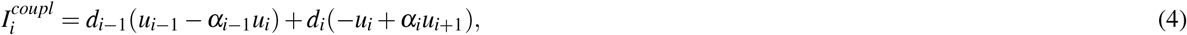

where 0 < *α_i_* ≤ 2 is the coefficient of asymmetry applied to the right-hand side cell in each coupled pair of model cells.

The cells playing the role of natural primary (SN) and subsidiary (SP7 through SP10, FP8, and PB) pacemakers are gray-shaded on the scheme (Fig 2). They are described by the same APm with a negative coefficient *a*_1_. The oscillating (pacemaking) properties of the APm were considered recently^28^. Implementing the oscillating APm cells simplifies the model description, providing easy control of intrinsic oscillation frequency and reliable overdrive suppression^29^. That substitutes a significant advantage of the cardiac APm over the FitzHugh-Nagumo and modified Van der Pol models and their combinations, used in the modeling of the simplified cardiac conduction system^13, 30^, due to the absence of the quiescent state in the modified Van der Pol model and the absence of well-defined threshold potential in both FitzHugh-Nagumo and modified Van der Pol models. Also, close values of the minimal diastolic potential of both non-pacemaking and pacemaking APm cells allow the driving of a relatively big number of non-pacemaking cells by a single pacemaking APm cell^28^.

### 1.2 Parameters

As a starting point for the search of optimal model cell parameters, we took the experimental data from^2, 31^: minimal and maximal AVN conduction time for normal case and after the FP and SP ablations for anterograde conduction and retrograde conduction, the effective refractory period for control case and after the SP ablation, for both anterograde conduction and retrograde conduction.

For the sake of simplicity, in the course of the model development, we paid attention to the differences between the FP and SP conduction delays but not the conduction velocities. Faster conduction velocity in the FP is not confirmed in experiments^32^. Anatomically shorter FP distance compared with the SP is considered a primary cause of a shorter conduction delay in the FP^33^. Thus, though in our phenomenological model, the SP is longer than the FP in terms of the number of model cells (10 vs. 8), this difference in the model cell numbers may not correspond to the real anatomical structure of the rabbit AVN. Proper pathway conduction delays were obtained by setting different distributions of the coupling coefficient *d* and coupling asymmetry α values and refractory periods of the model cells (Fig 3). In this regard, in our model, the cells represent groups of real cells and are distinguished not by biophysical types^3, 10, 34^ but by a gradual change of the parameters. Fig 3 shows the distribution of refractoriness in the uncoupled model cells. The refractory period values are in line with the available experimental data: 166 ±30 ms for the rabbit SN and 68 ±11 ms for atrial muscle^35^, 91 ±10 ms for the inferior nodal extension^31^, and 141 ±15 ms for the transitional zone^36, 37^, which is an anatomical substrate of the FP.

**Figure 3.**
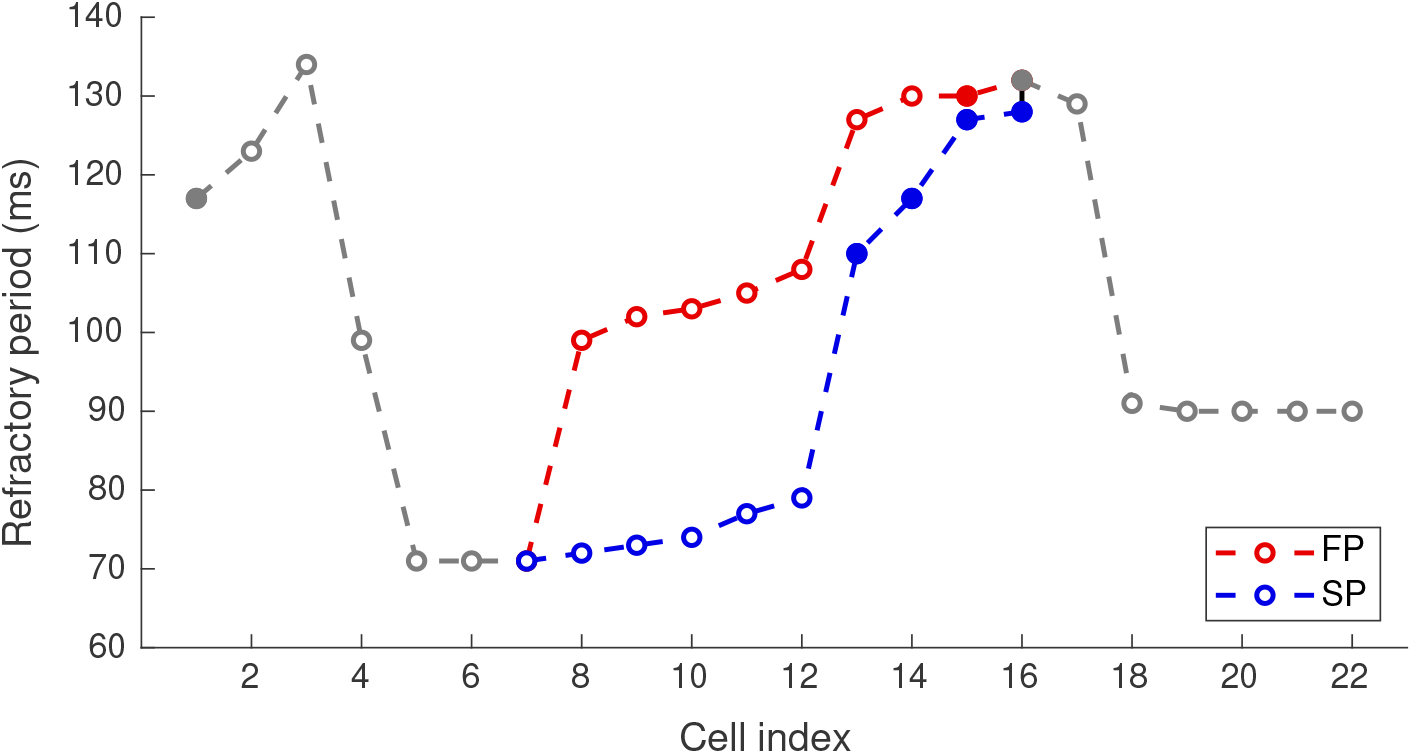
Distribution of refractory period for uncoupled model cells. Opaque and filled circles denote excitable and pacemaking cells, respectively.

The FP and SP ablations were simulated by setting coupling coefficient *d* = 0 in the middle of the pathways (between FP5 and FP6, and SP6 and SP7 cells).

### 1.3 External Pacing

The S1S2 stimulation protocol consists of ten basic S1 impulses followed by a premature S2 impulse. We created the AVN conduction curves applying constant S1S1 basic cycles with 360 ms intervals (shorter than spontaneous sinus rhythm of 361 ms) followed by the test premature S1S2 cycle with stepwise reduction starting from the basic cycle length until the full AVN functional refractory block occurs (360–90 ms)^2, 37^. The current stimulation impulses were of 1 ms duration and 1.3 × greater than the threshold value (*I^stim^* = 280.0). The AFB and AFL were simulated by applying similar stimulation impulses through the intermediate cell AM* to eliminate the influence of the pacing impulse shape. The correct shape of AM* action potentials allowed more natural stimulation of the atrial part of the conduction system model. For the simulation of the AFB, a random sequence of the impulses with the same amplitude and duration was generated using MATLAB pseudorandom number generator with uniform distribution in the 75–125 ms range.

### 1.4 Visualization of Laddergrams

An original algorithm was developed to visualize the laddergrams with the exact activation timing of each model cell. It allows tracing the wavefront propagation for each excitation sequence and in each pathway separately (with red color for the FP, and blue color for the SP) based on the activation time in each cell. The activation times in each model cell were recorded at the level of 0.25 during the action potential upstroke.

For anterograde propagation, the algorithm determines the leading pathway which provides the excitation to reach the PB first, thus, the traces coming down to the HB are marked in the same color. This is very useful since it allows visual differentiation between conduction through the SP and FP obtained in the simulations and easy comparison with the experimental method^11, 38, 39^, in which pronounced change in the signal amplitude in the inferior HB domain occurs when conduction switches from one pathway to another, and is especially important for visualization of chaotic rhythms^40^.

### 1.5 Numerical Methods

The solution of the set of Eqs 1-2 was performed with MATLAB (R2022a) standard ODE solver *ode23* with relative tolerance 10^−7^ and absolute tolerance 10^−10^. The *ode23* is a single-step solver and implements the explicit Runge-Kutta (2,3) pair^41^. Since *ode23* uses an adaptive time stepping, for the reliable periodic and random pacing, MATLAB *mod* function was used to force the solver to accept input impulses at predefined time moments. The solver output with the variable time steps was evaluated with the finite discretization of 0.1 ms for further analysis. Open-end boundary conditions were used at both SN and HB6 ends of the model.

## 2 Results and their validation

### 2.1 Remarks on Figures of Excitation Propagation

Figs 4–8 demonstrate different cases of simulated excitation propagation in the AVN. Each figure consists of a scheme and three panels. The scheme illustrates the propagation conduction sequence of interacting pathways in each case. The top panel shows the laddergram with exact timing of each model cell excitation. The trace color in the HB part corresponds to the leading pathway. The middle and bottom panels demonstrate the sequences of action potentials passing between the SN and HB through the FP and SP, respectively.

**Figure 4.**
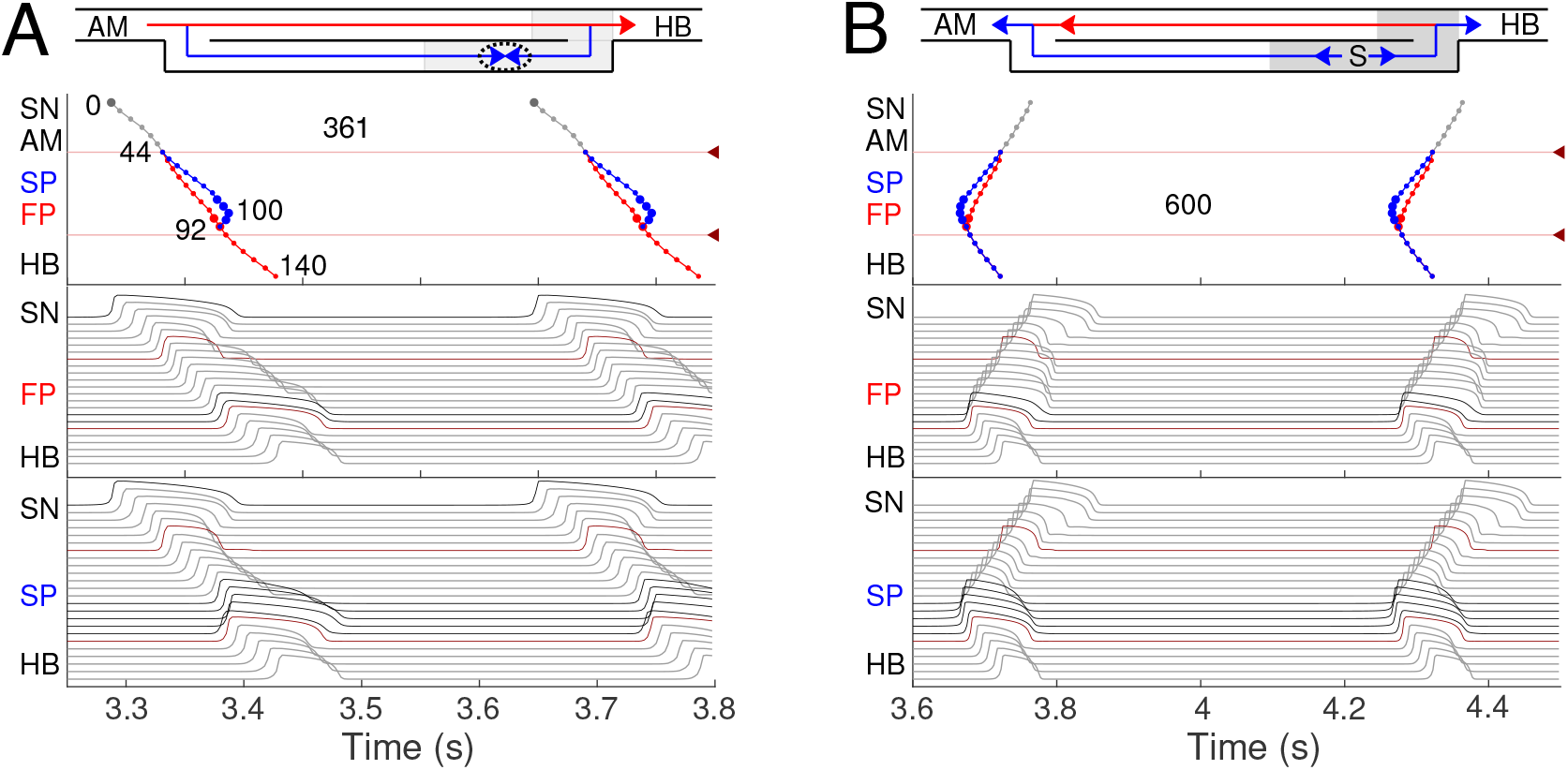
Primary and subsidiary pacemaking. Here and after *top scheme* illustrates propagation conduction sequences of interacting fast (FP, red arrows) and slow (SP, blue arrows) pathways. *Top panel*: the laddergram (Lewis ladder diagram) with the exact timing of each model cell excitation. Smaller circles denote quiescent excitable model cells, and larger circles - natural pacemaker cells. Red and blue traces correspond to the propagation through the FP and SP, respectively. The trace color in the HB part corresponds to the leading pathway. Dark red arrowhead markers on the right side and the adjacent lines denote the points of latency time measurements. *Middle panel*: the sequence of action potentials passing through the FP between the SN and HB. *Bottom panel:* the sequence of action potentials passing through the SP between the SN and HB. Action potentials in red correspond to the latency time measurement points, and in black color - to the pacemaking cells. Time scale is common for all three panels. **(A)** Rabbit’s normal sinus rhythm of 166 bpm (361 ms interval). The numbers on the traces indicate excitation wave latency with respect to the SN in milliseconds. **(B)** In the case of a silent SN pacemaker, the spontaneous activity with 100 bpm rate (600 ms interval) originates in the SP.

**Figure 5.**
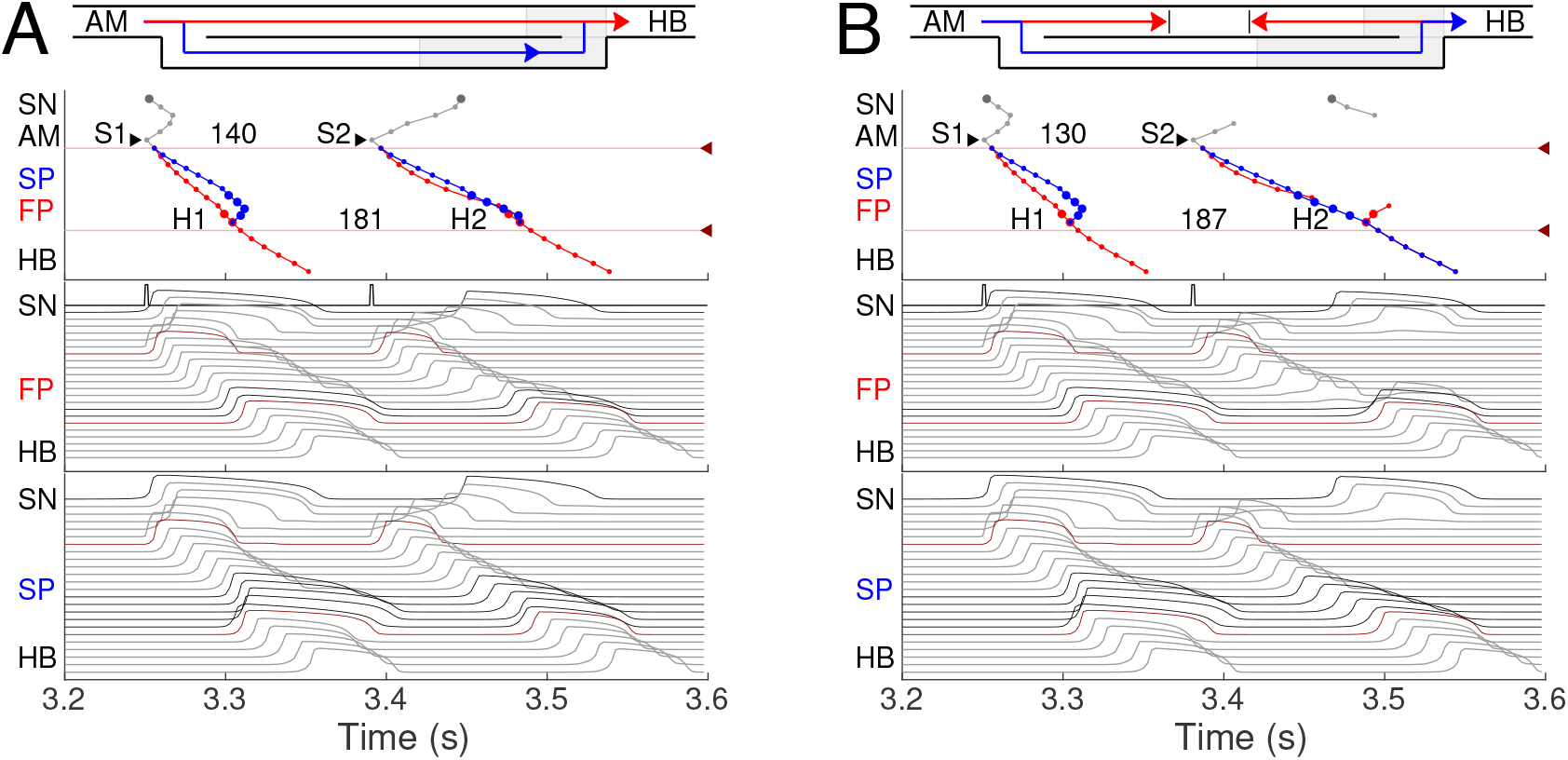
Atrial pacing in the control case with S1S2 interval of 140 ms and 130 ms. **(A)** At 140 ms interval, the excitation wave arrives via the FP and SP to the PB almost simultaneously with a slight advance of the FP. **(B)** The conduction in the FP is stopped due to a functional conduction block. In the top panel, arrowhead markers indicate points of stimulation. Upper (between the markers) and lower numbers correspond to the S1S2 and H1H2 intervals, respectively. Short stimulation impulses are shown at the top of the middle panel.

**Figure 6.**
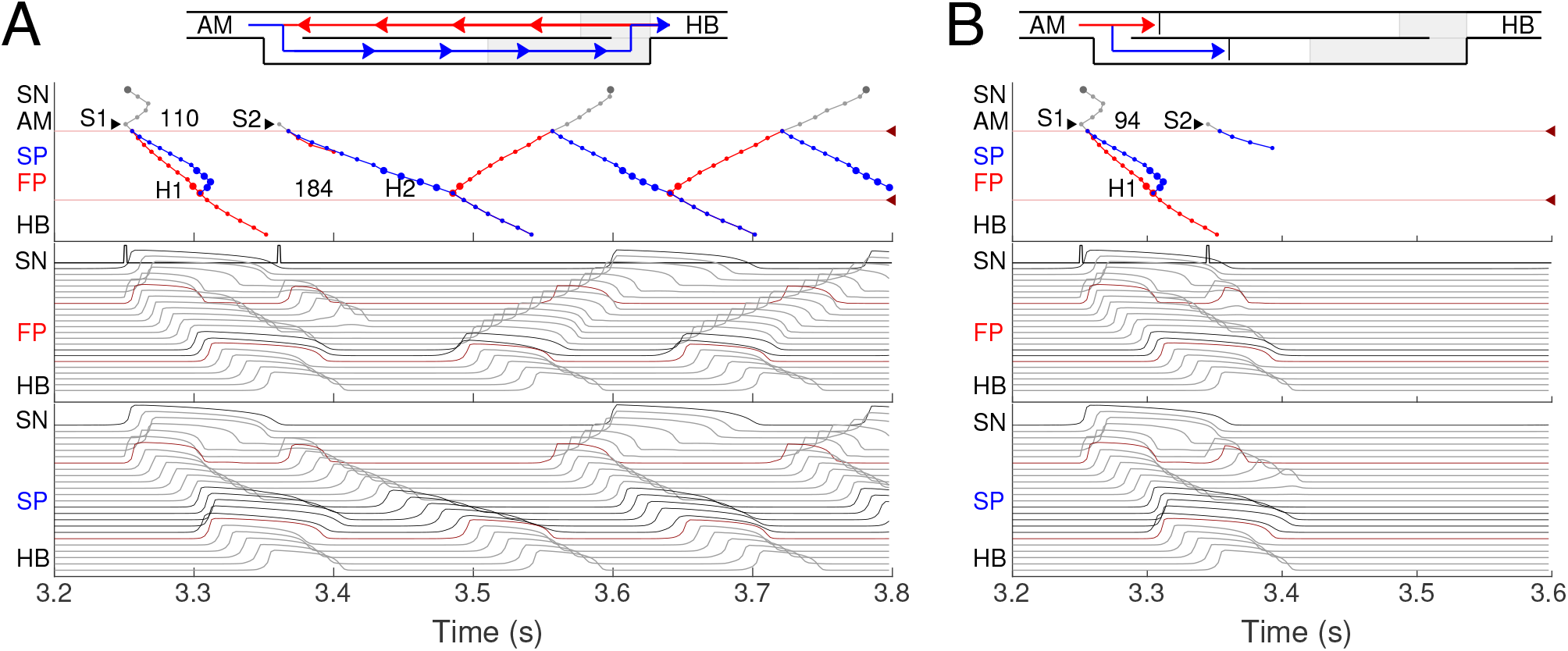
**(A)** Atrial pacing in the control case with S1S2 interval of 110 ms. Perpetual SP–FP AVNRT. **(B)**Atrial pacing for the control case with S1S2 interval of 94 ms. Full nodal conduction block.

**Figure 7.**
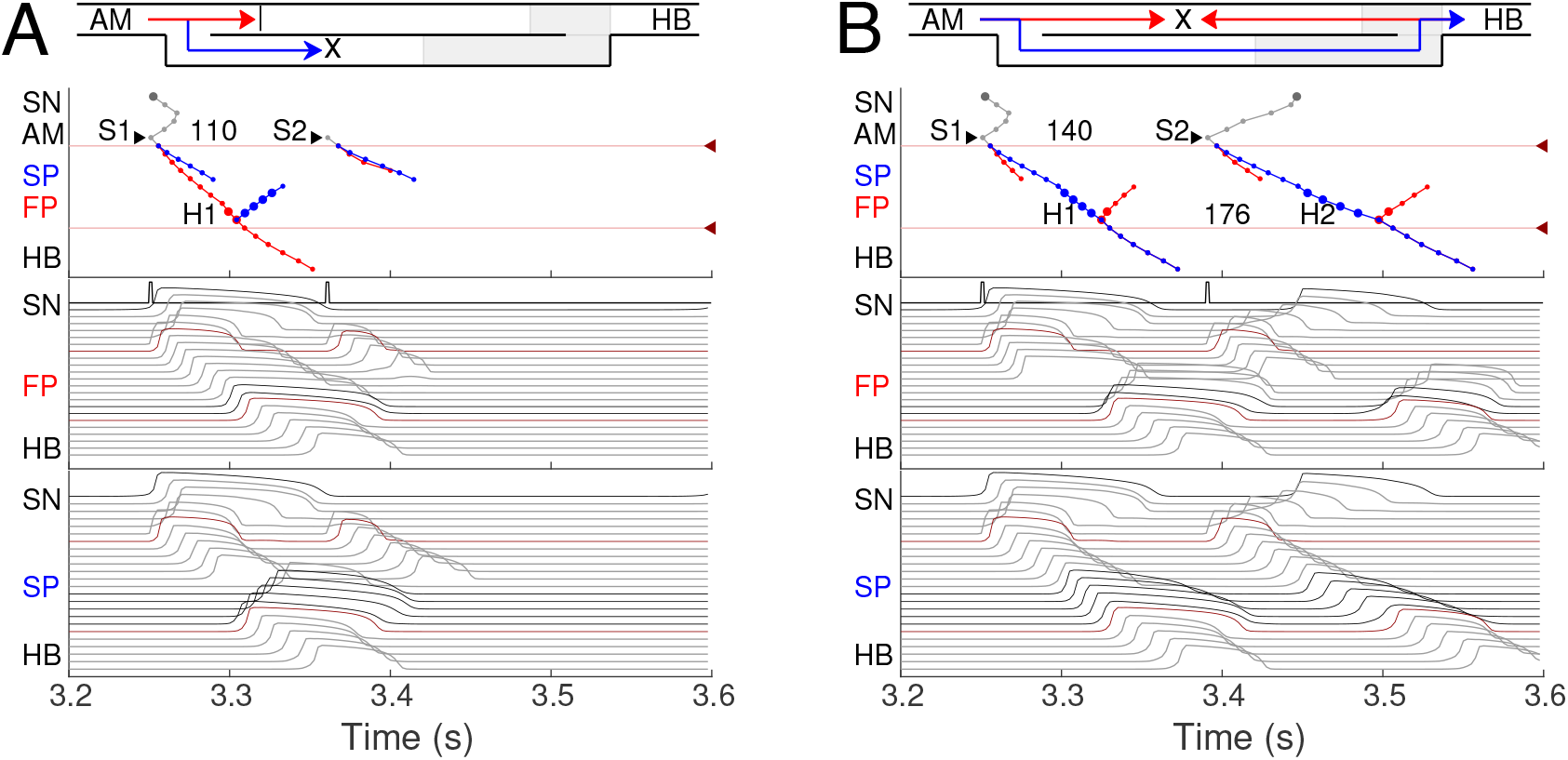
SP and FP ablations. **(A)**Atrial pacing with S1S2 interval of 110 ms after SP ablation. See Fig 6A for comparison. **(B)** Atrial pacing with S1S2 interval of 140 ms after FP ablation. See Fig 5A for comparison.

**Figure 8.**
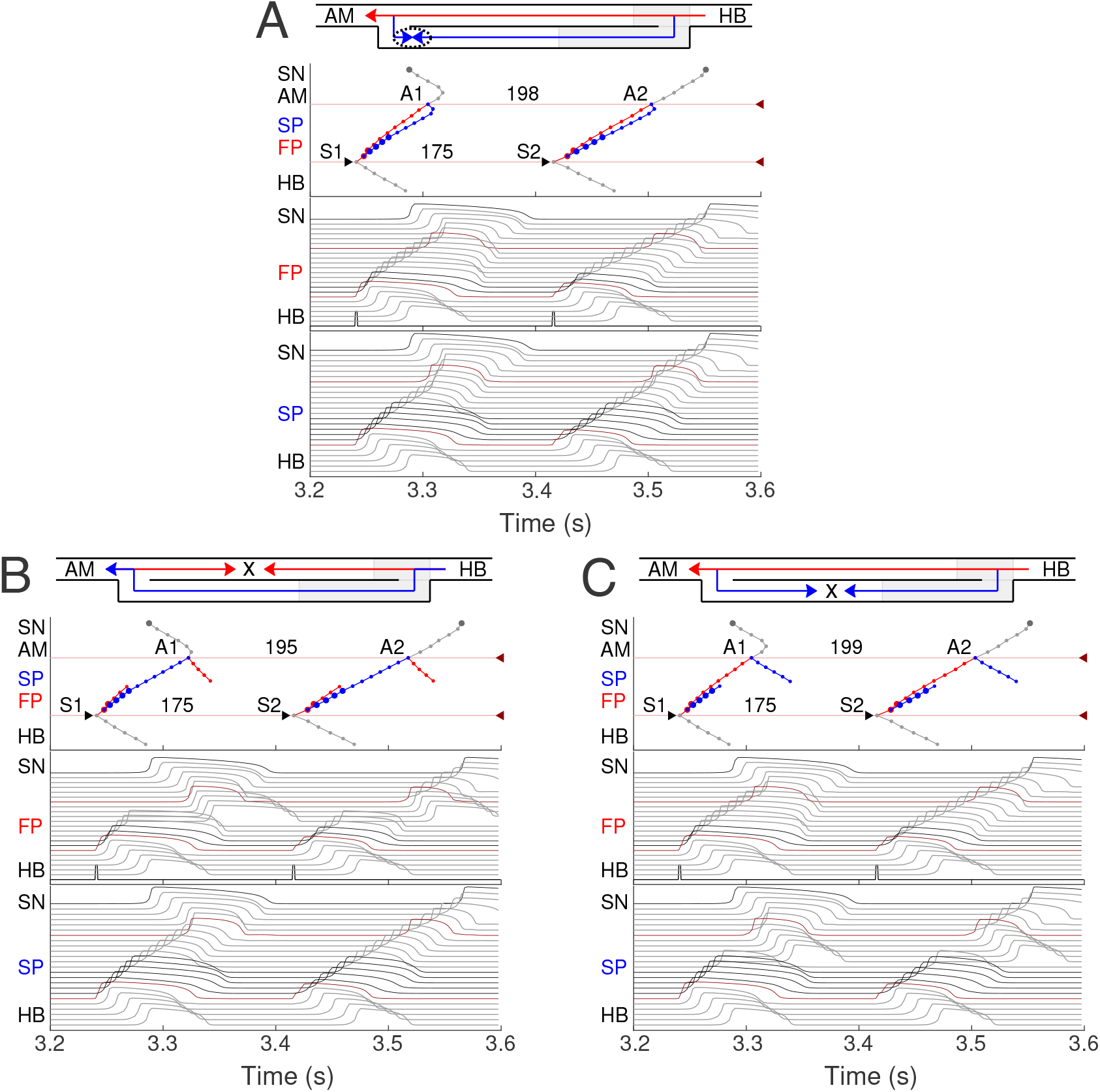
Retrograde AVN conduction, HB pacing with 175 ms S1S2 interval. **(A)** Control case. **(B)** After FP ablation. **(C)** After SP ablation. Upper and lower numbers correspond to A1A2 and S1S2 (H1H2) intervals. Short stimulation impulses are shown at the bottom of the middle panel.

### 2.2 Normal Sinus Rhythm

First, normal sinus rhythm with a cycle length of 361 ms (166 bpm) which is typical for rabbit heart (363 ±21 ms,^42^) was simulated, and the results are shown in Fig 4A. The numbers in the top panel indicate excitation wave latency with respect to the SN and the sinus rhythm in milliseconds. The excitation wave initiated at the SN propagates along the PS and AM cells and reaches the last atrium point (AM3) at 43.6 ms. Then the wave propagates through both the FP and SP. The wave in the FP arrives first at the PB cell at 92.3 ms, causing retrograde propagation through the SP. Both SP anterograde and retrograde excitation fronts meet in the inferior nodal extension at 99.8 ms and cancel each other. The leading FP conduction wave travels further down from the PB to the last HB cell of the model and arrives there at 139.6 ms. The obtained latency values are very close to the experimental measurements^3^ shown in Fig 1B for the right atrium of the rabbit heart and computer simulations with complex 1D ion-channel based rabbit AVN model^10^. A similar scheme of AVN conduction via the FP and SP was also observed in the case of atrial pacing at long S1S2 intervals (see Fig 5A below).

### 2.3 AVN Automaticity

In addition to its essential role in controlling the electrical impulse transmission between the atria and ventricles, the AVN can also serve as a subsidiary pacemaker when the SN fails^33^. In rabbits, the AVN leading pacemaker site is defined in the inferior nodal extension, generating electrical excitation with an average cycle length of about 600 ms (627±111 ms,^9^. In this case, the excitation spreads in two directions - upward to the atria and downward to the HB. Fig 4B shows the laddergram of AVN automaticity and the corresponding propagating action potentials in the absence of SN activity. The SP is the leading pathway for both anterograde and retrograde directions.

### 2.4 Atrial Pacing

The most important properties of the AVN behavior are reflected in the so-called AV nodal conduction or recovery curve^43, 44^. The characteristic steps of the AVN anterograde conduction curve calculation during atrial pacing are presented in Figs 5 and 6. In the range of long S1S2 intervals, the conduction scheme is similar to that demonstrated in Fig 4A with the FP leading.

At the S1S2 interval shortened to approximately 140 ms, a transitional stage occurs when the FP and SP excitation waves reach the PB almost simultaneously with minimal difference in latencies (Fig 5A). In such a case, while our model determines the leading pathway mathematically, from a physiological point of view both pathways provide conduction down to the HB. Simultaneous input of both pathways was experimentally demonstrated in HB electrograms with intermediate height^39, 40^.

For the S1S2 intervals shorter than 135 ms, only the SP becomes functional and allows passage of the excitation from the SN to the HB (Fig 5B).

By reducing the interval below 123 ms, we observed the onset of perpetual AVN reentrant tachycardia (AVNRT)^6, 8^, when excitation repeatedly circulates within the AVN ring (Fig 6A). The existence of AVNRT in the rabbit AVN at short S1S2 intervals was reported in previous experiments^36^. An excitation wave propagating anterogradely along the SP, retrogradely travels back up the FP, and then activates the atrial muscle, after which subsequent reentry cycles occur. This phenomenon takes place because the FP has a significantly longer refractory period than the SP (Fig 3). When excitation travels via the SP, the upper part of the FP is blocked, but the lower FP part recovers its excitability and conducts in a retrograde direction^2^. According to the AVNRT classification^8, 45^, the observed AVNRT belongs to the typical slow-fast type. AVNRT persisted with further shortening of S1S2 intervals until full functional block was achieved. In our simulations, the full functional block occurred at S1S2 intervals shorter than 95 ms when there was no conduction through both pathways (Fig 6B).

In all the above cases, the excitation wave from the SN pacemaker meets and annihilates with the retrograde wave from the atrial muscle caused by premature S1 impulses, and the SN activity is suppressed by S2 impulses.

### 2.5 Effects of FP and SP Ablations on Anterograde Conduction

Catheter ablation of the heart is used to destroy small areas in one of the AVN pathways, thereby stopping the propagation of unusual or undesirable electrical signals that travel through the AVN and cause irregular arrhythmias^46^. Fig 7 shows the simulation results of the atrial pacing after the SP and FP ablations. In the former case (Fig 7A), when S1S2 impulses are delivered at 110 ms intervals, the conduction of excitation is totally canceled since, at such short pacing intervals, functional conduction block in the FP takes place. Therefore, due to the broken AVN ring, SP ablation eliminates AVNRT shown in Fig 6A^8^.

FP ablation (Fig 7B) leads to increased conduction time^2^. Although the response to the S2 impulse looks similar to that shown in Fig 5B, where FP conduction is absent due to the functional conduction block associated with FP refractoriness, the increase in the conduction time is due to a different conduction propagation scenario caused by the preceding S1 impulse. Because ablation destroys pathway tissue, this conduction scenario extends over the entire range of S1S2 intervals (see complete anterograde conduction curves in Fig 9).

**Figure 9.**
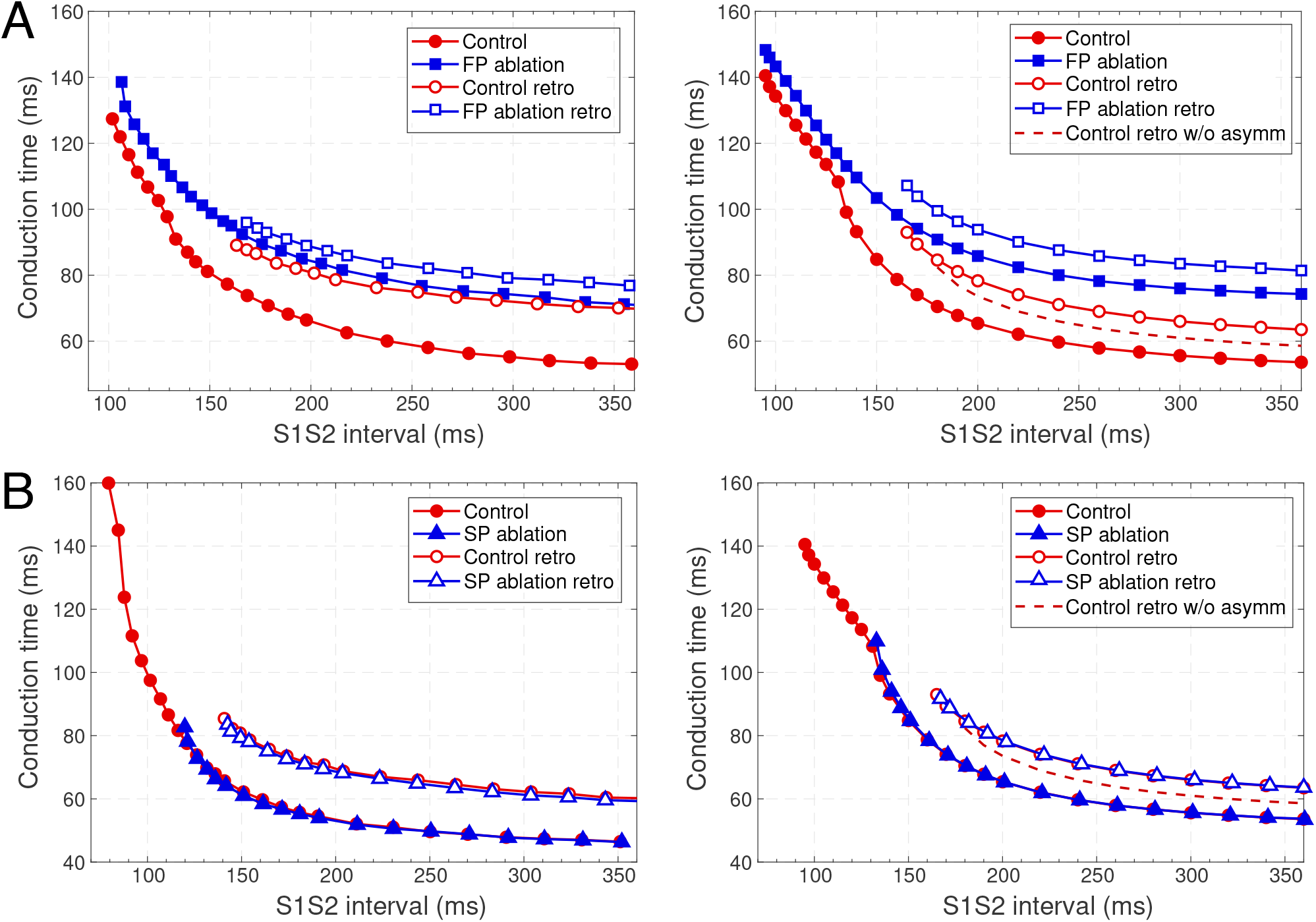
Comparison of experimental^31^ (left panels) and simulated (right panels) anterograde and retrograde recovery curves in the control case and after FP and SP ablations. **(A)** Experimental recovery curves in the control case and after FP ablation and simulated recovery curves. **(B)** Experimental recovery curves in the control case and after SP ablation and simulated recovery curves. Dashed lines correspond to the simulated retrograde conduction in the control case without coupling asymmetry.

### 2.6 Retrograde Conduction and FP and SP Ablations

Retrograde conduction refers to conduction from the lower part of the cardiac conduction system to the atria through the AVN. Experimental and clinical studies have demonstrated the ability of the AV node to conduct impulses in the anterograde and retrograde directions^2, 47–50^.

We simulated retrograde conduction applying S1S2 (H1H2) stimulation to the HB (at HB1 model cell, *Retro* arrow in the Fig 2) and plotted pertinent recovery curves (see AVN conduction curves Subsection). Fig 8 demonstrates retrograde conduction in the control case (Fig 8A), after FP ablation (Fig 8B), and after SP ablation (Fig 8C) for S1S2 interval of 175 ms.

In the control and SP ablation cases, excitation propagates retrogradely via the FP, creating almost the same A1A2 intervals and with close H1A1 and H2A2 conduction times. In the former case, similar to the case of atrial pacing with long intervals, retrograde and anterograde waves in SP meet and annihilate, but in the upper part of the AVN model (Fig 8A). In the FP ablation case, the A1A2 interval was shorter (Fig 8B), while conduction passing through SP leads to longer H1A1 and H2A2 latencies (see complete retrograde conduction curves in Fig 9).

### 2.7 AVN Conduction Curves

Applying the S1S2 stimulation protocol to the atrial and HB cells, we constructed various conduction curves for control (pre-ablation) and post-ablation cases. The recovery curve represents the atrial-His conduction time A2H2 for anterograde conduction or His-atrial conduction time H2A2 for retrograde conduction plotted either versus the S1S2 interval (A1A2 for atrial pacing and H1H2 for HB pacing) or versus the recovery interval (H1A2 for atrial pacing and A1H2 for HB pacing)^51^.

According to the experimental findings, the AVN conduction curves demonstrate pronounced direction-dependent behavior^2, 31^. Fig 9 shows the comparison of experimentally obtained (left panels) (modified from^31^) and simulated (right panels) anterograde and retrograde recovery curves in the control and ablation cases plotted versus S1S2 intervals. The modification from the dependence on recovery time H1A2 to S1S2 (A1A2) interval was performed by shifting each conduction curve on its minimal conduction time (A1H1) value according to the formula A1A2 = H1A2 + A1H1^51, 52^.

Simulated anterograde recovery curve (Fig 9A, right panel) in the control case demonstrates a typical exponential-like rise with reducing pacing interval^11, 12, 31^ with a pronounced bend and tilting at the point where conduction switches from the FP to SP with smooth change of the conduction curve from flat to steep without gap, which is typical for rabbit AVN^2, 11, 31^. At long S1S2 intervals, the SP conduction is overcome by the FP conduction (flat portion of the control curve, see also Fig 5A), while at short S1S2 intervals, excitation propagates through the SP only (Figs 5B and 6A) due to longer refractory period of the FP (Fig 3). In the S1S2 range of 95–122 ms, the perpetual AVNRT was observed^36^, as mentioned above (Fig 6A).

The nodal effective refractory period (ERPN) is set as the longest S1S2 pacing interval that results in neither HB nor atrial response in the cases of atrial pacing (anterograde propagation) and HB pacing (retrograde propagation), respectively. FP ablation markedly increased conduction time in both anterograde^31, 36^ and retrograde^31^ conduction cases. The ERPN was 94 ms for anterograde conduction and did not change after FP ablation (91±10 ms in^31^). For the retrograde conduction, the ERPN was 163 ms and also did not change after FP ablation (143±16 ms before and 149±14 ms after FP ablation in^31^).

During our initial simulations, the value of the upward shift of the retrograde control curve was relatively small (dashed line in Fig 9), while the experiments^31^ demonstrated much slower retrograde conduction in the control case. The introduction of the asymmetry to the intercellular coupling in the FP, PB, and upper part of HB allowed additional slowing of the retrograde control conduction, better reflecting the AVN direction-dependent conduction properties^2^.

SP ablation blocked its conduction, while FP conduction remained unaffected. The ablation led to the prolongation of ERPN to 132 ms (141±15 ms in^31^) and the disappearance of the steep part of the anterograde conduction curve in the short range of S1–S2 intervals (Fig 9B). The ablation also eliminated the AVNRT, as observed in clinical practice^8^ (compare Figs 6A and 7A).

In the retrograde case, SP ablation did not affect retrograde conduction and SP post-ablation curve virtually coincides with the control curve (Fig 9B). This, together with conduction scenarios shown in Figs 8A and 8C, confirms retrograde conduction through the FP in the control case. The ERPN was also prolonged to approximately the same value (163 ms) as in the retrograde pre-ablation case, matching with the experimental results (151 ±12 ms in^31^).

Fig 10 shows three main anterograde AVN functional curves (refractory curve H1H2, His-atrial curve H1A2, and recovery curve A2H2) from experiments (left panels) and our simulations (right panels). The AVN functional refractory period (FRPN) is determined as the shortest interval between two consecutive impulses propagated from the atria to the HB (H1H2). In our simulations, FRPN was 180.5 ms, which is in agreement with experiments (175±10 ms in^53^.

**Figure 10.**
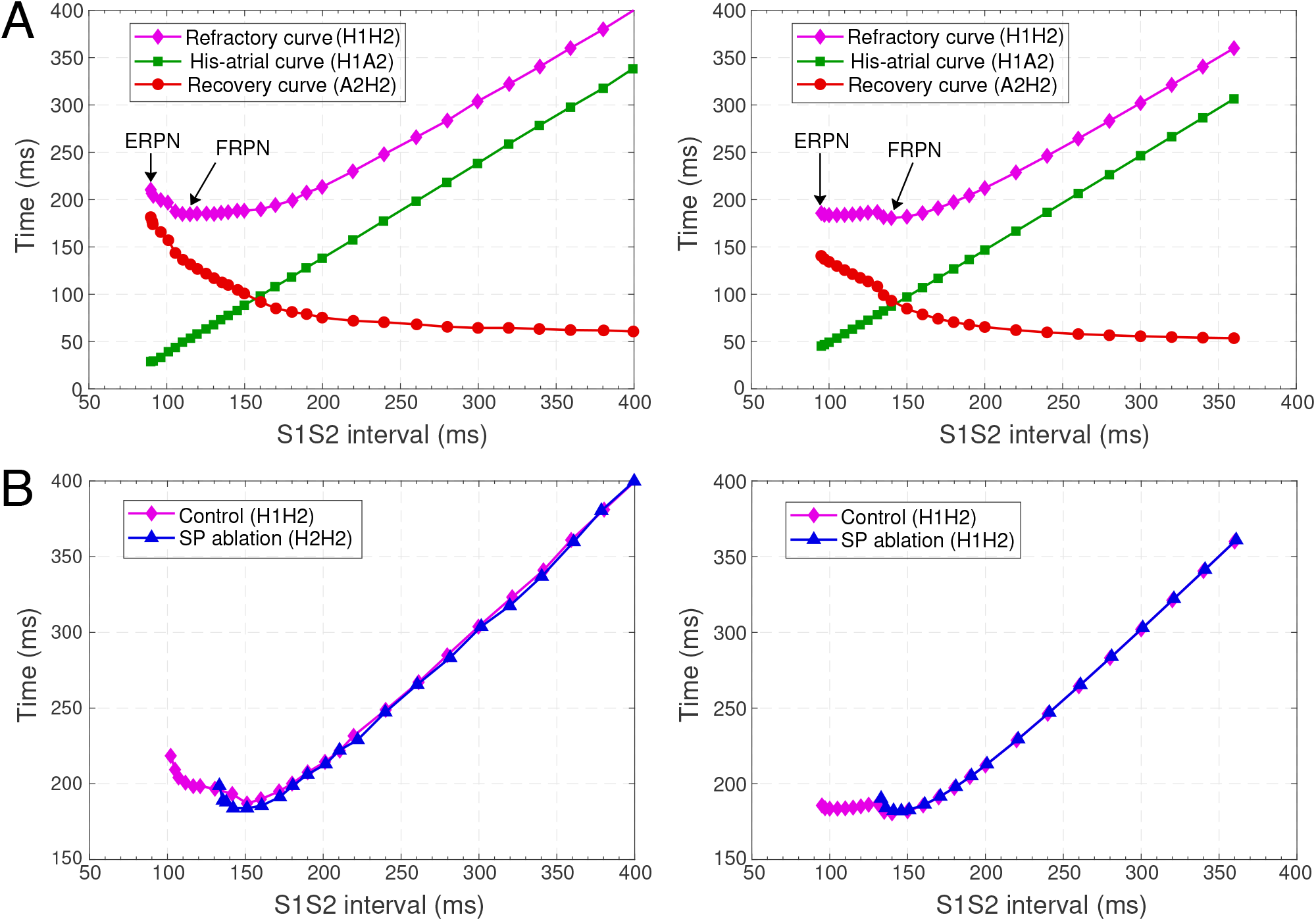
Comparison of experimental (left panels) and simulated (right panels) conduction curves. **(A)** Experimental^52^ and simulated AVN refractory, His-atrial, and recovery curves. **(B)** Experimental^54^ and simulated refractory curves before and after SP ablation. The simulated control refractory curves (H1H2) in the right panels are the same.

Each point of the refractory curve H1H2 obtained in our simulations equals the sum of A2H2 value on the recovery curve and H1A2 value on the His-atrial curve at the corresponding S1S2 interval (Fig 10A, right), which is in line with the previously proposed quantitative relationship between the three function curves^52^ (Fig 10A, left). Fig 10B demonstrates another comparison of experimental^54^ and simulated AVN refractory curves before and after SP ablation. SP ablation decreased the left steeply rising part of the refractory curve compared to the control curve, resulting in an increase in ERPN, but not FRPN.

Note that the experimental conduction curves shown in Figs 9 and 10 were obtained for different rabbit preparations and slightly different experimental setups^31, 52, 54^. In particular, the AVN conduction measurements before and after FP and SP ablations were performed separately for each group of rabbits, thus making the experimental conduction curves rather different in Figs 9 and 10. On the other hand, our model parameters were adjusted for the rabbit data corresponding to Fig 9A, thus our results show a greater quantitative difference with the experimental conduction curves demonstrated in Figs 9B and 10.

### 2.8 Filtering of Atrial Rhythm During Atrial Fibrillation

It has been demonstrated that dual pathways are involved in AVN conduction during AFB by limiting the number of atrial impulses transmitted to the ventricles^39^. To show the AVN filtering function and reveal the interaction between the pathways during AFB we set random atrial pacing with uniform distribution in the 75–125 ms range (Fig 11). In Fig 11, different conduction sequences for each atrial impulse are present, including complete passage of the excitation to HB through either pathway with the domination of the SP conduction over the FP (which may depend on mean random atrial rate), and simultaneous conduction block in both pathways. One can also see many episodes of retrograde propagation annihilating with an anterograde impulse resulting from a subsequent atrial stimulus. Besides, the sinus rhythm was completely suppressed by faster atrial pacing. This filtering behavior during AFB corresponds to the results observed in experiments^39, 55^ and in simulations with complex 1D ion-channel based model^10, 56^.

**Figure 11.**
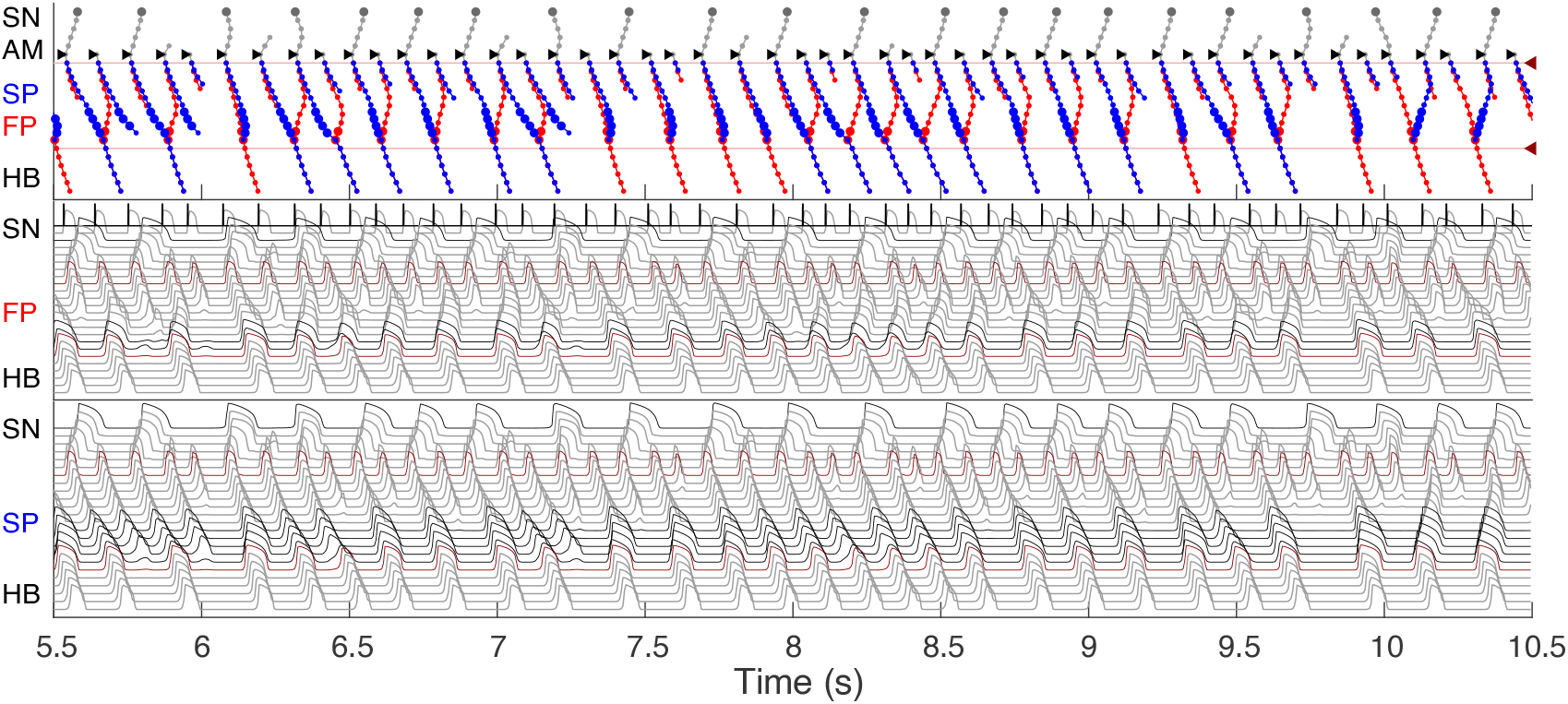
Filtering of atrial rhythm during atrial fibrillation stimulated by random impulse sequence in 75–125 ms range applied to AM* model cell. The short stimulation impulses and resulting AM* cell action potentials are shown in the top part of the middle panel.

As an additional validation of the AVN model filtering function, we simulated the effect of SP and FP ablations on the AFB. Fig 12A shows laddergrams of SP ablation (top panel) and FP ablation (bottom panel) cases simulated with the same random atrial impulse pattern, as shown in Fig 11. The HB rate was visually lower for the SP ablation case. This is also confirmed by the statistical distributions obtained for all three cases simulated with random atrial stimulation during 350 s (Fig 12B). The reduction of the mean HB rate from 339 bpm in the control case to 292 bpm in the SP ablation case and to 318 bpm in the FP ablation case and prolongation of the mean HH interval from 177 ms to 205 ms and 189 ms, respectively, revealed SP ablation as a relatively more effective way of ventricular rate control than FP ablation, which is in agreement with experimental^55^ and clinical^57^ findings, and with mathematical modeling study^11^.

**Figure 12.**
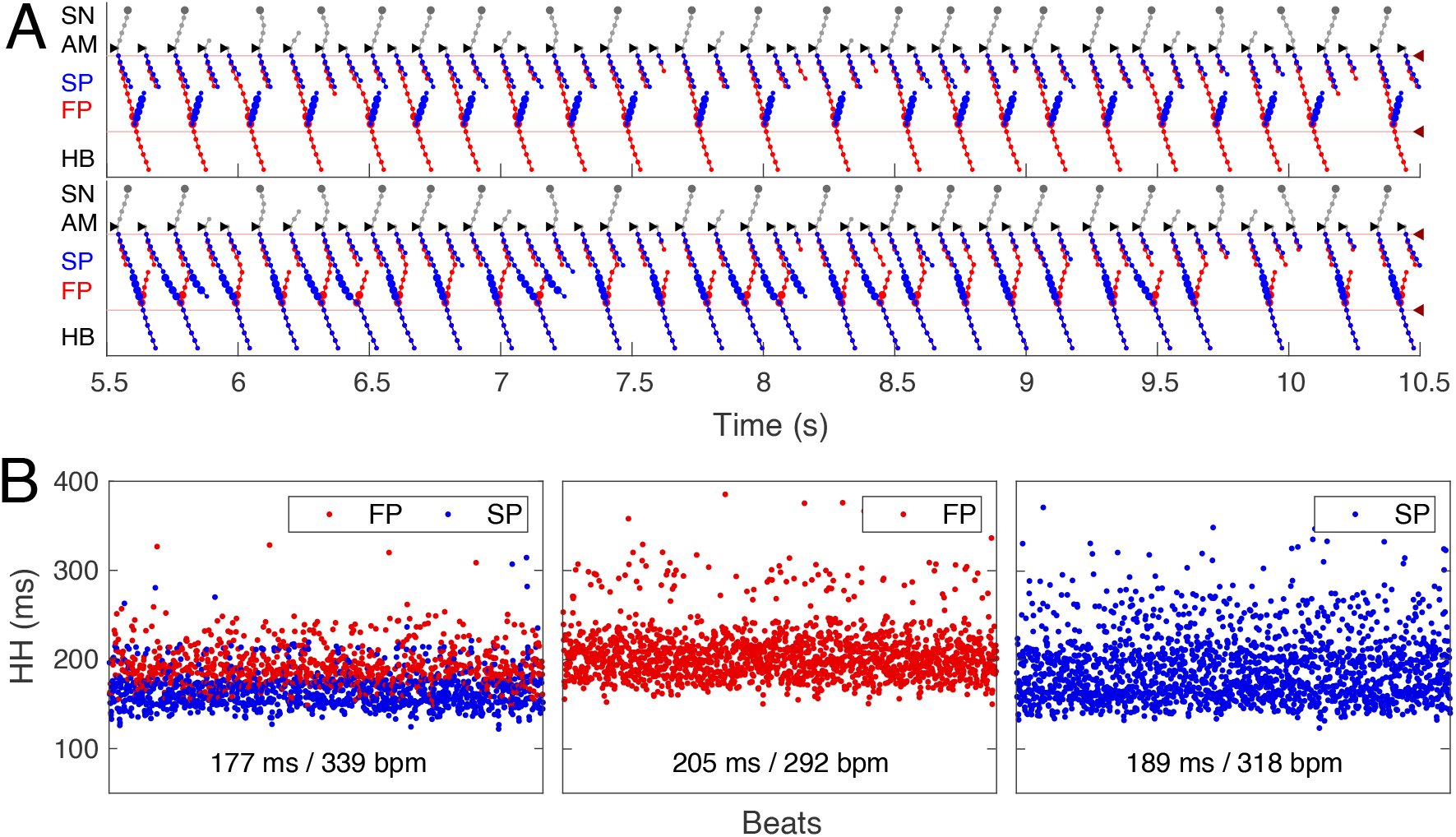
Impact of ablation on AVN filtering function. **(A)** Laddergrams of atrial fibrillation stimulated by the same random impulse sequence as in Fig 11 after SP ablation (top panel) and after FP ablation (bottom panel). **(B)** Distribution of consecutive HH intervals for intact AVN and after ablations. Random atrial pacing in the 75–125 ms range was applied for 350 s. Control case (left panel), after SP ablation (middle panel), and after FP ablation (right panel). Mean HH intervals and HB rates for each case are shown at the bottom.

### 2.9 Wenckebach Periodicity During Atrial Flutter

Dual pathway electrophysiology was shown to be also directly related to the occurrence of the Wenckebach phenomenol^11, 58^. Fig 13 demonstrates an example of regular atrial pacing mimicking AFL that yields 5:4 Wenckebach periodicity corresponding to four out of five atrial impulses conducted to HB, obtained at a regular atrial pacing interval of 123.0 ms (atrial/His rates 488/389 bpm). Fig 13A shows laddergram and action potential propagation through the FP and SP, while Fig 13B demonstrates AH conduction times (top panel) and HH intervals (bottom panel) of consecutive beats in the Wenckebach cycle. This “typical” Wenckebach pattern is characterized by the progressive increase of AVN conduction times (AH) and progressive shortening of HH intervals before the conduction block occurs^59^. Also, exactly in line with the experimentally obtained HB electrograms, the first atrial stimulus after the conduction block passed through the FP (depicted by red squares in Fig 13B), followed by a transition to the SP until its ultimate block^11, 58^.

**Figure 13.**
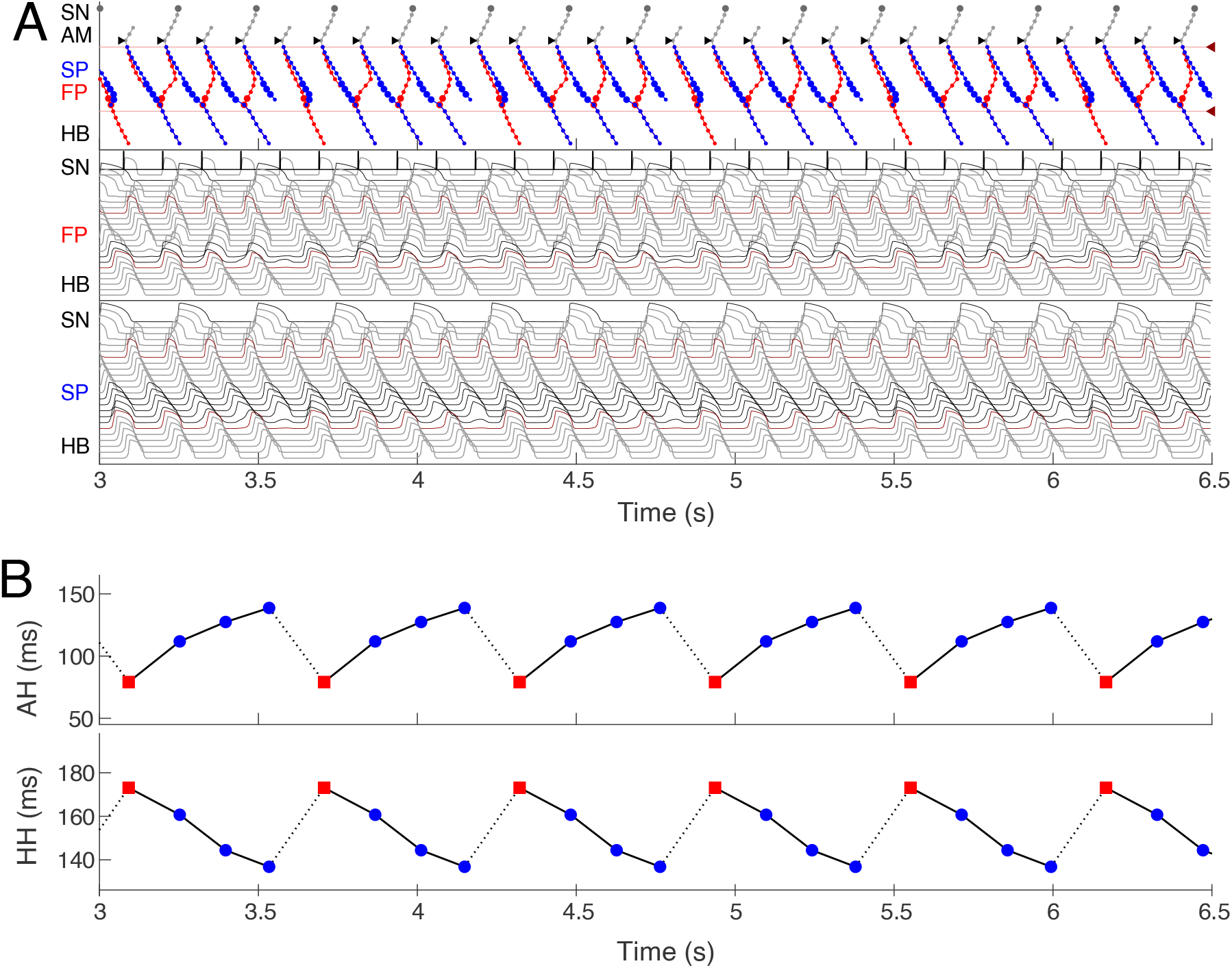
Simulated 5:4 Wenckebach pattern (atrial/His rates 488/389 bpm) at regular atrial pacing (atrial flutter) with the interval of 123.0 ms. **(A)** Laddergram and action potential propagation through the FP and SP. **(B)** AH conduction time of consecutive beats in the Wenckebach cycle (top panel), and HH interval (bottom panel). Each Wenckebach cycle begins with FP conduction (red squares), AH delays progressively prolonged, and HH intervals progressively shortened.

## 3 Discussion

### 3.1 Major Achievements

In this work, we have developed a compact one-dimensional mathematical model of rabbit AVN extended structure, which includes SN and HB. The AVN model is based on the Aliev-Panfilov nonlinear system of two-variable differential equations, describing both quiescent excitable and pacemaking cells. Despite its simplicity and a small number of model cells (33 elements), the model is multi-functional and does not require structural changes for reproduction of a wide variety of AVN dual-pathway electrophysiological phenomena. In particular, the model can correctly reproduce normal sinus rhythm, AVN subsidiary pacemaking, AVNRT, filtering AVN function during AFB and AFL with Wenckebach patterns, and situations after FP and SP ablations.

For the first time, we simulated the retrograde conduction in the AVN and implemented the asymmetry of the intercellular coupling in the FP, PB, and upper part of HB for accurate reproduction of AVN direction-dependent conduction properties^2^. Several conduction curves (Figs 9 and 10) reflecting the general behavior of AVN were presented along with experimental data to demonstrate their good agreement.

Our model is accompanied by a visualization tool, which discloses processes taking place inside the AVN “black box” by creating laddergrams and revealing the leading pathway for anterograde propagation. With the help of the laddergrams, it became possible to compare FP and SP functional state before and after ablation, including hidden SP activation over a broad range of atrial and HB pacing intervals, as well as the involvement of dual AVN pathways in the Wenckebach phenomena and during AFB.

We believe the AVN model is suitable for cardiac research and educational purposes. It can be integrated into the 3D atrial^14^ or whole-heart models^60, 61^ for near-realtime or even real-time simulations. The parameters of the proposed model can be adjusted for other animal species. Another possible application of the model with some modifications according to human AVN physiology is utilization in the testbed systems for implantable cardiac devices^62, 63^. The model can also be used for quick preliminary testing of hypotheses before accurate and detailed simulations with computationally expensive and complex ion-channel cellular models.

### 3.2 Comparison with Previous AVN Models

Similar to the complex multi-cellular ion-channel model^10^ and functional mathematical model^11^, our model is based on experimental rabbit data. Due to the small number of elements and much simpler nonlinear equation systems, our model allows faster computation and easier tuning than more biophysically detailed model^10^, keeping the model structure unchanged. On the other hand, functional mathematical models like^11^ benefit from effortless fitting of exponential equations to particular rabbit cardiac preparation. The presented model is capable of reproducing almost all electrophysiological properties of AVN found in both types of AVN models^10, 11^, with the additional feature of retrograde conduction with realistic ERPN in control and FP and SP post-ablation cases.

### 3.3 Limitations

The concept of the presented model is based on a 1D cable structure where FP and SP are electrically isolated from each other. Thus, possible effects related to the complexity of real 3D AVN structure are not taken into account. For example, the source-sink (current-to-load) mismatch^64^, when a relatively small bunch of cells (the source) provides insufficient total current to pass the excitation to a larger inactive cell group (the sink), or expression of connexins in gap junctions^65–67^, can be the reason for the difference in conduction velocity in anterograde and retrograde directions, reflected in the direction-dependent conduction curves (Fig 9). This also can explain the quantitative difference between experimental and simulated anterograde conduction curves in the range of short S1S2 pacing cycles and the retrograde conduction cases.

Due to the simplicity of the two-variable APm used in this study, the proposed AVN model does not allow simulation of subtle effects on cardiac cell ion channels, such as the impact of drugs or change in ion concentrations as demonstrated in the more complex multicellular ion-channel based models^10, 56^.

## 4 Conclusion

In this work, we have developed a compact multi-functional model of a rabbit atrioventricular node with dual pathways. The model is relatively simple, contains a small number of model cells, and is computationally efficient, allowing for fast simulations. Using the developed model, we successfully reproduced several electrophysiological phenomena accompanied by visualization of laddergrams. We believe the proposed AVN model has a high potential for utilization in research and education both as an independent module and as a part of complex three-dimensional atrial or whole-heart simulation systems. The proposed model can also be used for quick preliminary testing of hypotheses before accurate and detailed simulations with computationally expensive and complex cellular models.

## Data Availability Statement

Data supporting the finding of this study are available from the corresponding author upon reasonable request.

## Author Contributions

MR wrote the program code and performed computer simulations. ER analyzed the results. All authors contributed to the conception and design of the study and manuscript preparation; read and approved the submitted version.

## Acknowledgments

The authors would like to thank Katrina Armstrong for her assistance with the manuscript.

## Supplemental Data

**Supplementary movie S1.**Animated S1S2 anterograde stimulation in the control case.

**Supplementary movie S2.**Animated S1S2 retrograde stimulation in the control case.

